# Modeling the SARS-CoV-2 nsp1–5’-UTR complex via extended ensemble simulations

**DOI:** 10.1101/2021.02.24.432807

**Authors:** Shun Sakuraba, Xie Qilin, Kota Kasahara, Junichi Iwakiri, Hidetoshi Kono

## Abstract

Nonstructural protein 1 (nsp1) of severe acute respiratory syndrome coronavirus 2 (SARS-CoV-2) is a 180-residue protein that blocks translation of host mRNAs in SARS-CoV-2-infected cells. Although it is known that SARS-CoV-2’s own RNA evades nsp1’s host translation shutoff, the molecular mechanism underlying the evasion was poorly understood. We performed an extended ensemble molecular dynamics simulation to investigate the mechanism of the viral RNA evasion. Simulation results showed that the stem loop structure of the SARS-CoV-2 RNA 5’-untranslated region (SL1) is recognized by both nsp1’s globular region and intrinsically disordered region. The recognition presumably enables selective translation of viral RNAs. Cluster analysis of the binding mode and detailed analysis of the binding poses revealed several residues involved in the SL1 recognition mechanism. The simulation results imply that the nsp1 C-terminal helices are lifted from the 40*S* ribosome upon the binding of SL1 to nsp1, unblocking translation of the viral RNA.

## Introduction

SARS-CoV-2 (severe acute respiratory syndrome coronavirus 2) belongs to *Betacoronaviridae*, and is the causative pathogen of COVID-19. Nonstructural protein 1 (nsp1) resides at the beginning of SARS-CoV-2’s genome, and it is the first protein translated upon SARS-CoV-2 infection. After self-cleavage of open reading frame 1a (orf1a) by an orf1a-encoded protease (nsp3; PLpro), nsp1 is released as a 180-residue protein. SARS-CoV-2 nsp1 is homologous to nsp1 of SARS-CoV-1, the causative pathogen of SARS, sharing 84 % sequence identity with the SARS-CoV-1 protein. Nsp1 functions to suppress host gene expression^1–6^ and induce host mRNA cleavage,^1,2,7,8^ effectively blocking translation of host mRNAs. The translation shutoff hinders the host cell’s innate immune response icluding interferon-dependent signaling. ^1,9^ Multiple groups have recently reported cryogenic electron microscopy (cryo-EM) structures of SARS-CoV-2 nsp1–40*S* ribosome complexes. ^10–12^ The structural analysis showed that two *α*-helices are formed in the C-terminal region (153-160, 166-179) of nsp1 and binds to the 40*S* ribosome. These helices block host translation by shutting the ribosomal tunnel used by the mRNA. This blockade inhibits formation of the 48*S* ribosome pre-initiation complex, which is essential for translation initiaion. ^3,12^ But while nsp1 shuts down host mRNA translation, it is nown that the viral RNAs are translated even in the presence of the nsp1, and that they evade degradation.^2–4^

These mechanisms force infected cells to produce only viral proteins instead of normal host cell proteins; indeed, in a transcriptome analysis, 65 % of total RNA reads from Vero cells infected with SARS-CoV-2 were mapped to the viral genome.^13^ It has also been shown that nsp1 recognizes the 5’-untranslated region (5’-UTR) of the viral RNA^4,6,11^ and selectively enables translation of RNAs that have a specific sequence. The first stem loop in the 5’-UTR^4,6,14^ has been shown to be necessary for translation initiation in the presence of nsp1. Specifically, with SARS-CoV-1,^4^ bases 1-36 of the 5’-UTR enable translation of viral RNA; with SARS-CoV-2, bases 1-33^14^ or 1-40^6^ of the 5’-UTR of SARS-CoV-2 enable translation. However, the precise molecular mechanism remains poorly understood.

In the present research, therefore, our aim was to describe the detailed mechanism by which SARS-CoV-2 RNA evades nsp1. We modeled and simulated a complex complsed of SARS-CoV-2 nsp1 and the SARS-CoV-2 5’-UTR’s first stem loop using an extended ensemble molecular simulation. A detailed analysis of the simulation suggests the molecular basis of nsp1 recognition of the 5’-UTR, whereby interaction of nsp1 and the stem loop prevents nsp1’s C-terminal helices from binding to the ribosomal tunnel.

## Methods

### Simulation setup

We constructed a complex of nsp1 and 5’-UTR of SARS-CoV-2 RNA. Nsp1 is a partially disordered 180-residue protein, in which the structures of residues 12-127 and 14-125 have been solved by X-ray crystallography in SARS-CoV and SARS-CoV-2, respectively. The structures of other residues (1-11, 128-180) are unknown, and residues 130-180 are thought to be an intrinsically disordered region (IDR).^15,16^ We constructed the SARS-CoV-2 nsp1 structure using homology modeling based on the SARS-CoV-1 nsp1 conformation (Protein Data Bank (PDB) ID: 2HSX^15^). Modeling was performed using MODELLER. ^17^ We noted that SARS-CoV-1 nsp1 and SARS-CoV-2 nsp1 are aligned without gaps. The structure of the IDR was constructed so as to form an extended structure. For nsp1, we used the AMBER ff14SB force field^18–21^ in the subsequent simulations.

The initial structure of the RNA stem was constructed using RNAcomposer.^22,23^ Bases numbered 1-35 from the SARS-CoV-2 reference genome (NCBI reference sequence ID NC_045512.2)^24^ were used in the present research. This sequence corresponds to the first stem loop of the SARS-CoV-2 RNA 5’-UTR. Hereafter, we will call this RNA “SL1.” SL1 was capped by 7-methyl guanosine triphosphate (m7G-ppp-). The first base (A1) after the cap was methylated at the 2’-O position to reflect the viral capped RNA. Charges and bonded force field parameters for these modified bases were respectively prepared using the restrained electrostatic potential (RESP) method^25^ and analogy to existing parameters. For SL1, we used a combination of AMBER99 + bsc0 + *χ*OL3.^18,19,26,27^ To maintain the structural stability of the stem loop, we employed distance restraints between the G-C bases. Specifically, between residues G7–C33, G8–C32, C15–G24 and C16–G23, distance restraints were applied such that the distances between the N1, O6 and N2 atoms of guanosine and the N3, N4 and O2 atoms of cytidine, did not exceed 4.0 Å. Between these atoms, flat-bottom potentials were applied, where each potential was zero when the distance between two atoms was less than 4.0 Å, and a harmonic restraint with a spring constant of 1 kJ mol^−1^ Å^−2^ was applied when it exceeds 4.0 Å. We used `acpype`^28^ to convert the AMBER force field files generated by AmberTools^29^ into GROMACS.

The nsp1 and SL1 models were then merged and, using TIP3P^30^ water model with Joung-Cheatham monovalent ion parameters^31^ (73,468 water molecules, 253 K ions, 209 Cl ions), were solvated in 150 mM KCl solution. The initial structure is presented in Fig. 1A. A periodic boundary condition using a rhombic dodecahedron unit cell was used with a size of *ca.* 140 Å along the X-axis. Note that we started the simulation from the unbound state; that is, nsp1 and SL1 were not directly in contact with each other. The total number of atoms in the system was 224,798.

**Figure 1:**
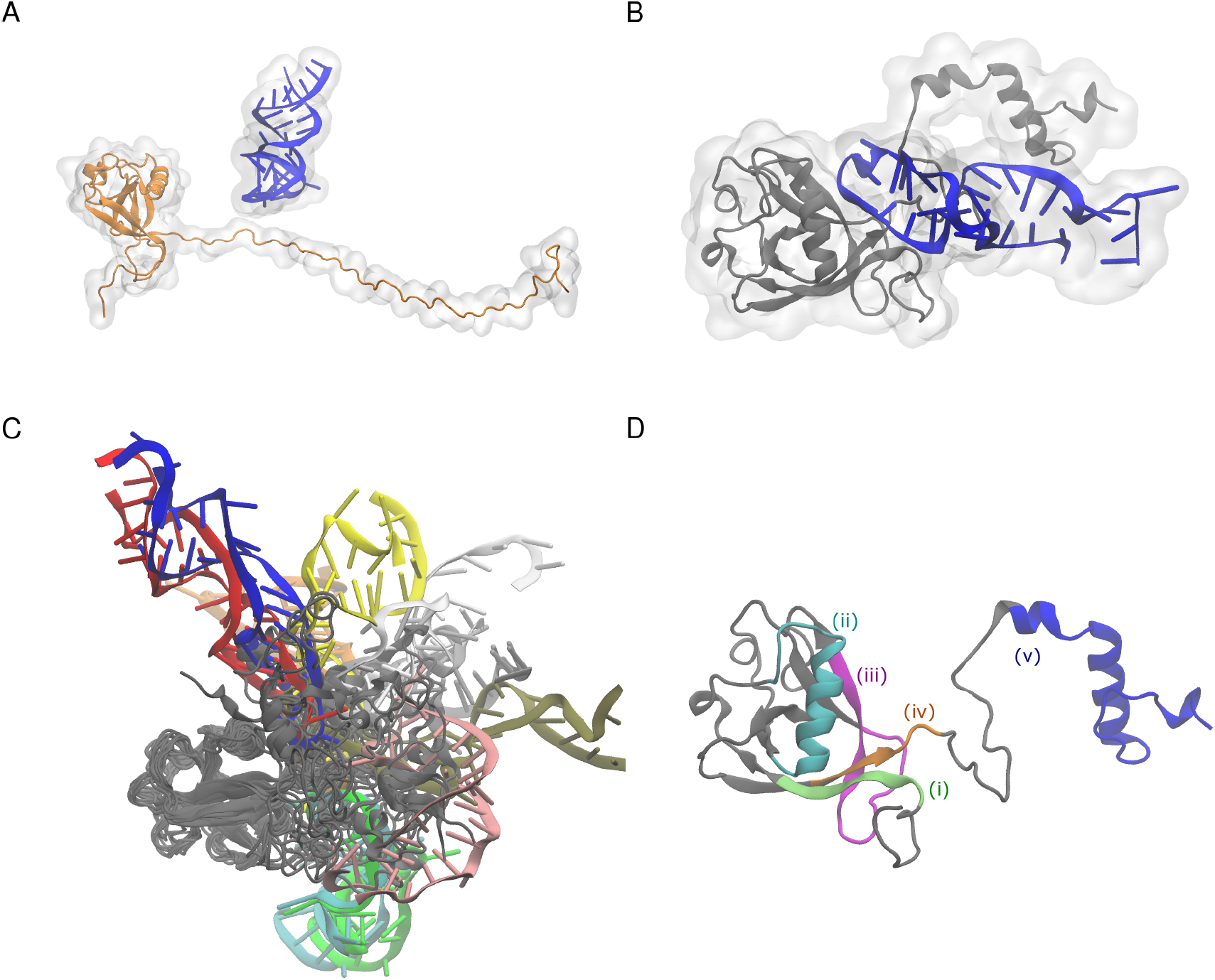
(A) Initial structure before starting the simulation. (B) Structure of the complex at 50 ns in the 0th replica (i.e., the simulation with the unscaled potential). (C) Structures from superimposition of 20 representative snapshots of the nsp1-SL1 complex. Snapshots were obtained from a weighted random sampling. Different snapshots from SL1 are colored differently. (D) Nsp1 segmentation used in the analysis: (i) residues 1 to 18, green; (ii) residues 31 to 50, cyan; (iii) residues 74 to 90, magenta; (iv) residues 121 to 146, orange; (v) residues 147 to 180, blue.

**Figure 2:**
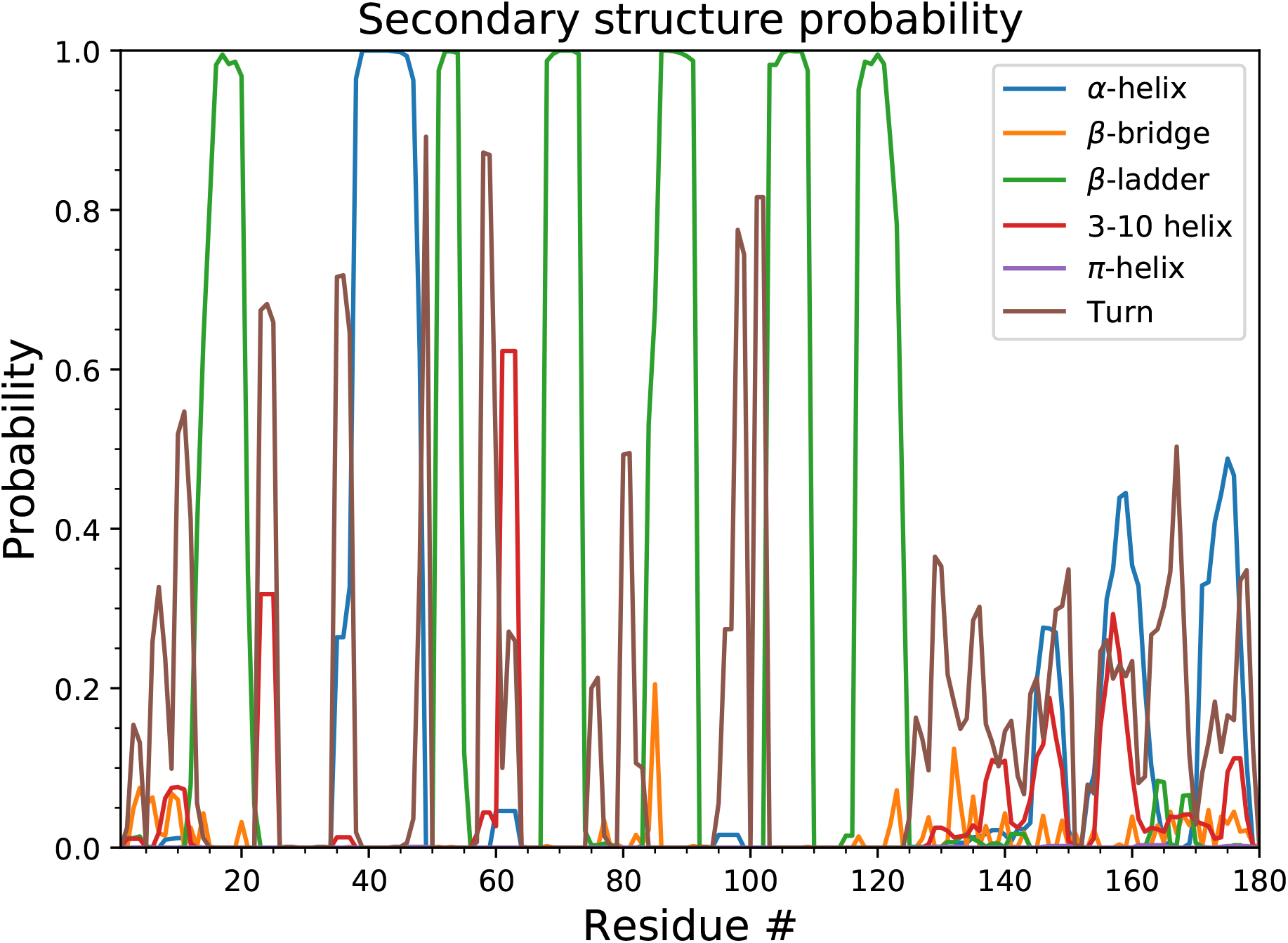
Secondary structure distribution of nsp1. Probabilities were calculated using the reweighting of the last 25 ns simulation trajectories.

Although it is possible to perform a molecular dynamics (MD) simulation of an nsp1-SL1 complex, due to the excessive charges on both molecules, the model tends to be trapped around the initial configuration of the complex in conventional MD simulations. Authors have previously shown that the sampling for nucleic acid–protein systems can be effectively solved by extended ensemble simulations. ^32–34^ In this work, we used replica exchange with solute tempering (REST) version 2 to sample various configurations of SL1 and the nsp1 IDR.^35^ We set both the disordered region (nsp1 1-11 and 128-180) and the entire SL1 as the “hot” region of the REST2 simulation. Note that in addition to the charge scaling for nsp1 and SL1, we also scaled the charges of counter-ions to prevent unneutralized system charge in the Ewald summation. The total number of replicas used in the simulation was 192. The replica numbered 0 corresponds to the simulation with the unscaled potential. In the final replica (numbered 191), nonbonded potentials between “hot”-“hot” groups were scaled by 0.25. Exchange ratios were 53-78 % across all replicas. To prevent numerical errors originating from the loss of significant digits, we used a double-precision version of GROMACS as the simulation software.^36^ We also modified GROMACS to enable the replica exchange simulation with an arbitrary Hamiltonian. ^37^ The patch representing modifications is supplied in the supporting information.

The simulation was performed for 50 ns (thus, 50 ns×192 = 9.6 μs in total), and the first 25 ns were discarded as an equilibration time. The simulation was performed with NVT and the temperature was set to 300 K. The temperature was controlled using the velocity rescaling method.^38^ The timestep was set to 2 fs, and hydrogens attached to heavy atoms were constrained with LINCS. ^39^

### Simulation analysis

To obtain the proper structural ensemble under the unmodified potential function, we used the multistate Bennett acceptance ratio (MBAR) method.^40,41^ With that method, we obtained a weighted ensemble corresponding to the canonical ensemble (trajectory with a weight assigned on each frame) from multiple simulations performed with different potentials. Only eight replicas corresponding to the eight lowest replica indices (i.e., the one with the unscaled potential function and seven replicas with the potentials closest to the unscaled potential) were used in the MBAR analysis. The weighted ensemble of the trajectory was used in the subsequent analyses. The ensemble of structures with the weight information is available online at https://bsma.pdbj.org/entry/26. Visualization was performed using VMD^42^ and pymol.^43^ The secondary structure of nsp1 was analyzed using the secondary structure definition of DSSP^44^ with `mdtraj`.^45^ We tested the convergence of the ensemble using the secondary structure distribution and the stability of the hydrogen bonds between nsp1 and SL1 (see the supplementary material for details).

### Interactions between nsp1-SL1

We applied three criteria to detect interactions between nsp1 and SL1. (i) Inter-residue contacts were detected with the criterion that the inter-atomic distance between the C*α* of an amino acid residue and C4’ of a nucleotide residue was less than or equal to 12 Å. (ii) Hydrogen bonds were detected with the criteria that the hydrogen-acceptor distance was less than 2.5 Å and the donor–hydrogen–acceptor angle was greater than 120 degrees. (iii) Salt-bridges were detected with the criterion that the distance between a phosphorous atom in the RNA backbone and the distal nitrogen atom of Arg or Lys was less than 4.0 Å.

### Clustering

On the basis of the inter-residue contact information, the binding modes of the nsp1–SL1 complex observed in the ensemble were evaluated by applying the clustering method. The inter-residue contact information in each snapshot was represented as a contact map consisting of a 180 × 36 binary matrix. The distance between two snapshots was then calculated as the Euclidian distance of vectors with 180 × 36 = 6480 elements. We applied the DBSCAN method^46^ to classify the binding modes. We arbitrarily determined two parameters, `eps` and `minPts`, for the DBSCAN method to obtain a reasonable number of clusters each of which had distinct binding modes. Note that the DBSCAN generates clusters each of which has more than `minPts` members based on the similarity threshold `eps`. The clusters with fewer than `minPts` members (including singletons) were treated as outliers. We used `eps` = 6 and `minPts` = 200 in this research. We also tested another clustering algorithm, OPTICS, ^47^ and confirmed that the two different methods generate qualitatively similar results (data not shown).

## Results and discussion

### The IDR partially forms secondary structure and binds to SL1

Although we did not restrain the RNA-nsp1 distance in the simulation and started the simulation with the two molecules apart, they formed a complex within the canonical ensemble. Figure 1 (B) shows a representative snapshot of the complex at the end of the simulation. The RNA stem binds to the C-terminal disordered region. However, as shown in Fig. 1 (C), when the N-terminal domain of nsp1 was superimposed, the RNA structures did not have a specific conformation. This implies that there was no distinct, rigid structure mediating nsp1-RNA binding.

We next investigated the secondary structure of the nsp1 region. Although we started the simulation from an extended configuration, the C-terminal region at residues 153-179 partially formed two *α*-helices, which is consistent with the fact that the C-terminal region forms two helices (residues 153-160, 166-179) and shuts down translation by capping the pore that mRNA goes through in the Cryo-EM structural analysis. The result also indicates that the cap structure may be formed before nsp1 binds to the ribosome, reflecting a pre-existing equilibrium, although the ratio of the helix-forming structures is only up to 50%. In addition to these known helices, residues 140-150 also weakly formed a mixture of *α*-helix and 3-10 helix. Residues at other regions (1-11, 128-139) remained disordered.

### Distance between the nsp1 N-terminal domain and C-terminal helices

Recent cryo-EM structures suggest that the N-terminal domain of nsp1 reside on the 40*S* ribosome, though the density map is ambigous (Fig. 3). Inspired by the structures, we investigated the geometric restraints on nsp1 in the presence of SL1. The distance between the center-of-mass of the nsp1 N-terminal domain (defined by residues 14-125) and that of the C-terminal helices (residues 153-179) was calculated and its histogram is plotted in Fig. 3(C). The distance distribution had two peaks at 27 and 33 Å, which is below 49.8 Å estimated from the cryo-EM structure (see supporting information for details). Indeed, 90.7% of the trajectory had a distance less than the experimentally estimated distance of 49.8 Å. This indicates that the configuration observed in the cryo-EM structure, which does not include SL1, is unlikely to happen when nsp1 is complexed with the SL1.

**Figure 3:**
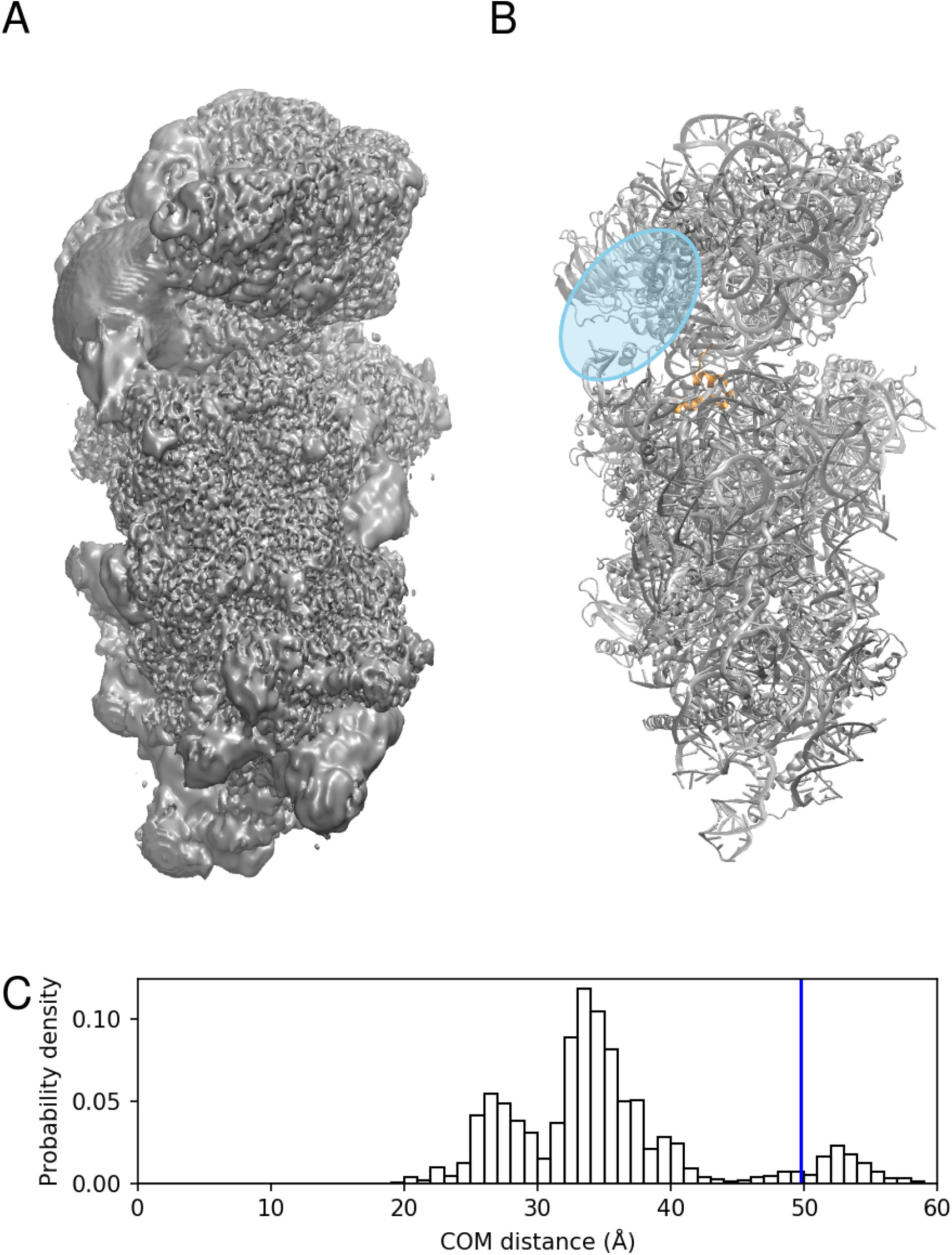
(A) Cryo-EM structure of nsp1-40*S* ribosome complex (Electron Microscopy Data Bank ID: EMD-11276). (B) Cartoon representation of the nsp1-40*S* ribosome complex (PDB ID: 6ZLW). The C-terminal helices of nsp1 are colored orange. The nsp1 N-terminal domain is thought to bind at the blue shaded region. (C) The distribution of the distance between the center-of-mass of the N-terminal domain and that of the C-terminal helices in our simulation. A blue vertical line indicates the distance estimated from the cryo-EM structure.

### SL1’s hairpin is recognized by the nsp1 IDR

Inter-residue contact probabilities between nsp1 and SL1 in the canonical ensemble are summarized in Table 1 and Fig. 4. Based on the distribution of the interactions, we categorized the binding interface of nsp1 into five regions (Fig. 1D and Supplementary Table S1): (i) the N-terminus (residues 1–18), (ii) the *α*1 helix (residues 31–50), (iii) the disordered loop between *β*3 and *β*4 (residues 74–90), (iv) the N-terminal side of the IDR (residues 121–146), and (v) the C-terminal side of the IDR (residues 147–180). These five regions interacted primarily with bases around C20 of the RNA fragments, which compose the stem loop. The most important region for recognition of SL1 was region (iv), the N-terminal side of the IDR. The probability of contacts between any residue in this region and SL1 was 97.4%. In particular, contact between Asn126 and U18 was observed in 84.1% of the canonical ensemble. The most frequently observed hydrogen bond in the canonical ensemble was Arg124–U18, the probability of which was 26.0% (Table 1). The second most important interface region was region (ii), *α*1 helix, which has two basic residues (Arg43 and Lys47), that frequently formed salt-bridges with the backbone of SL1. At least one salt-bridge in this region was included in 69.8% of the canonical ensemble. The third most important was region (iii), consisting of the loop between *β*3 and *β*4; 63.2 % of the canonical ensemble included at least one contact in this region. Asp75 sometimes formed hydrogen bonds with the bases of SL1. Regions (i) and (v) tended not to form hydrogen bonds or salt-bridges, but frequently contacted residues in these regions; the probability for interactions with regions (i) and (v) were 72.1% and 59.2%, respectively.

**Table 1:**
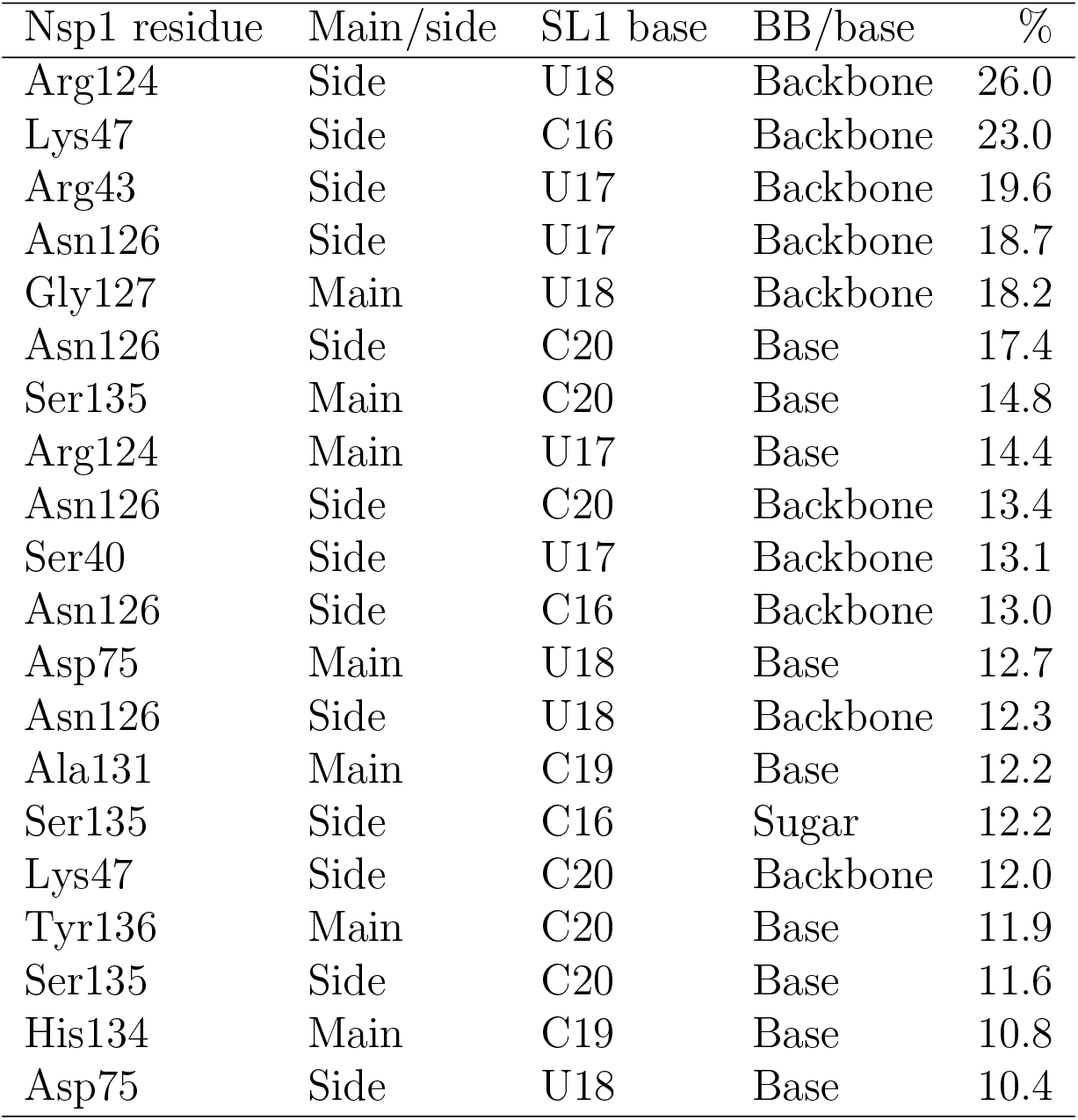
Hydrogen bonds observed between SL1 and nsp1.

| Nsp1 residue | Main/side | SL1 base | BB/base | % |
| --- | --- | --- | --- | --- |
| Arg124 | Side | U18 | Backbone | 26.0 |
| Lys47 | Side | C16 | Backbone | 23.0 |
| Arg43 | Side | U17 | Backbone | 19.6 |
| Asn126 | Side | U17 | Backbone | 18.7 |
| Gly127 | Main | U18 | Backbone | 18.2 |
| Asn126 | Side | C20 | Base | 17.4 |
| Ser135 | Main | C20 | Base | 14.8 |
| Arg124 | Main | U17 | Base | 14.4 |
| Asn126 | Side | C20 | Backbone | 13.4 |
| Ser40 | Side | U17 | Backbone | 13.1 |
| Asn126 | Side | C16 | Backbone | 13.0 |
| Asp75 | Main | U18 | Base | 12.7 |
| Asn126 | Side | U18 | Backbone | 12.3 |
| Ala131 | Main | C19 | Base | 12.2 |
| Ser135 | Side | C16 | Sugar | 12.2 |
| Lys47 | Side | C20 | Backbone | 12.0 |
| Tyr136 | Main | C20 | Base | 11.9 |
| Ser135 | Side | C20 | Base | 11.6 |
| His134 | Main | C19 | Base | 10.8 |
| Asp75 | Side | U18 | Base | 10.4 |

**Figure 4:**
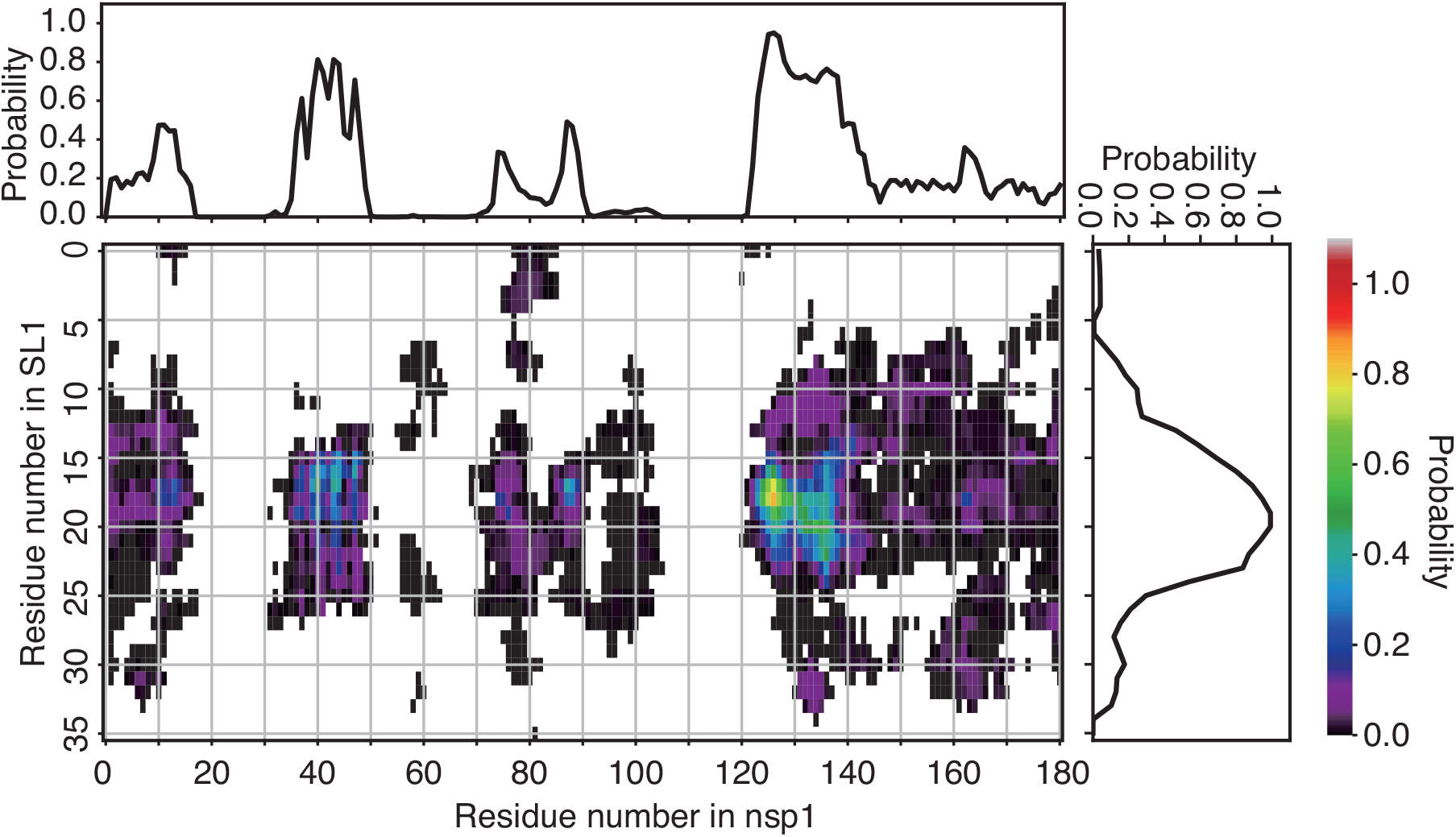
Residue-wise, all-against-all contact probability in the canonical ensemble. The color at each grid point indicates the statistical weight of the contact between the corresponding pair of residues (color scale is shown at the right of the panel). The points filled by white indicate no detectable probability of contacts. The line plots at the top and right of the contact map depict the contact probability for each residue, regardless of its counterpart.

**Figure 5:**
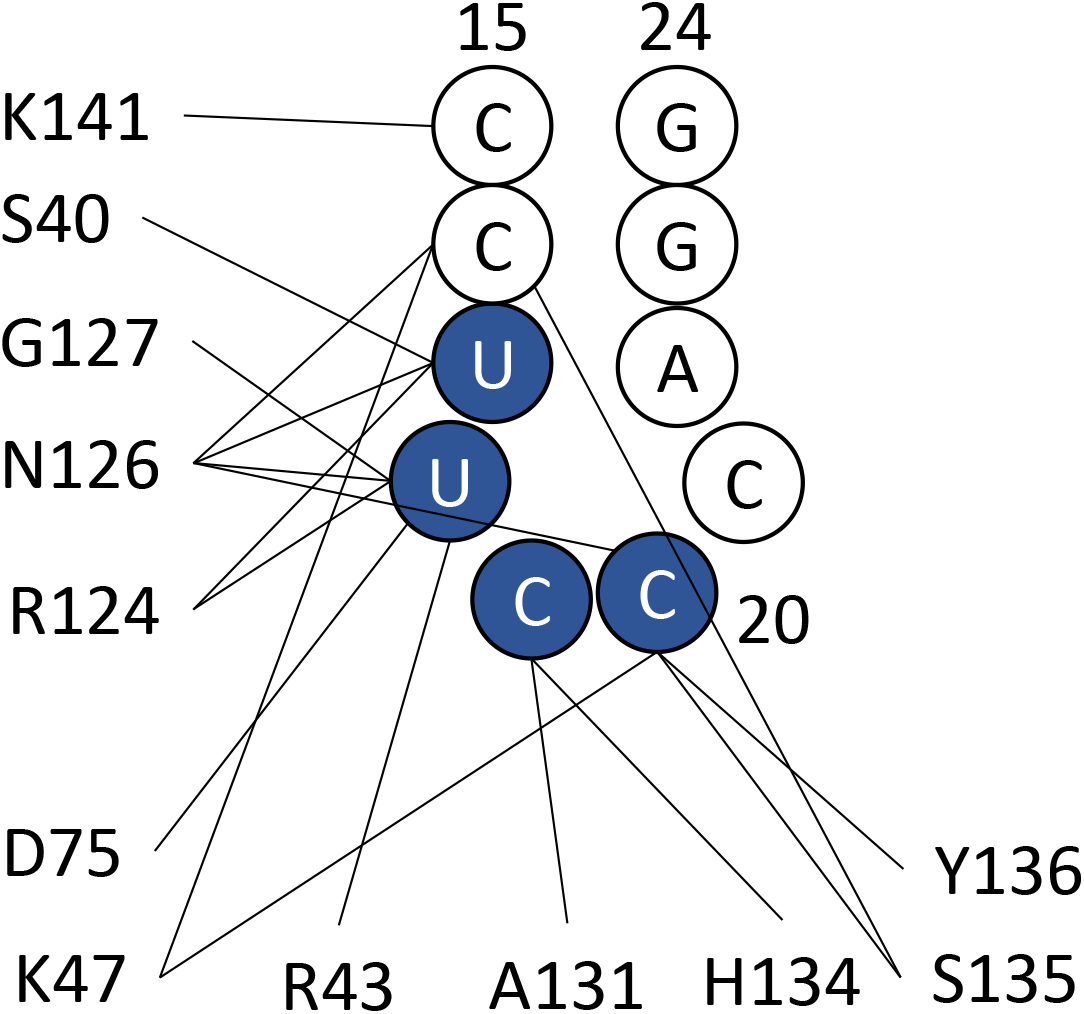
Graphical representation of the hydrogen bond interactions between SL1 and nsp1. Bases of U17 to C20 (colored blue) are recognized by the hydrogen bonds.

As an overall shape, the nsp1 surface consists of positive and negative electrostatic surface patches separated by a neutral region (Fig. 6A).^48^ The *α*1 helix in region (i) forms the interface between these two patches; one side of the helix contains basic residues (Arg43 and Lys47), and the other side contains some hydrophobic residues (Val38, Leu39, Ala42, and Leu46). The positive side of the *α*1 helix assumes a mound-like shape with a positively charged cliff (Fig. 6(B)). The bottom of the valley formed by the N-terminus and *β*3-*β*4 loop, or regions (i) and (iii), respectively, also contains positive electrostatic potentials. The positively charged cliff and valley attract and fit to the negatively charged backbone of SL1. Eventually the IDRs in region (iv) and (v) grab SL1.

**Figure 6:**
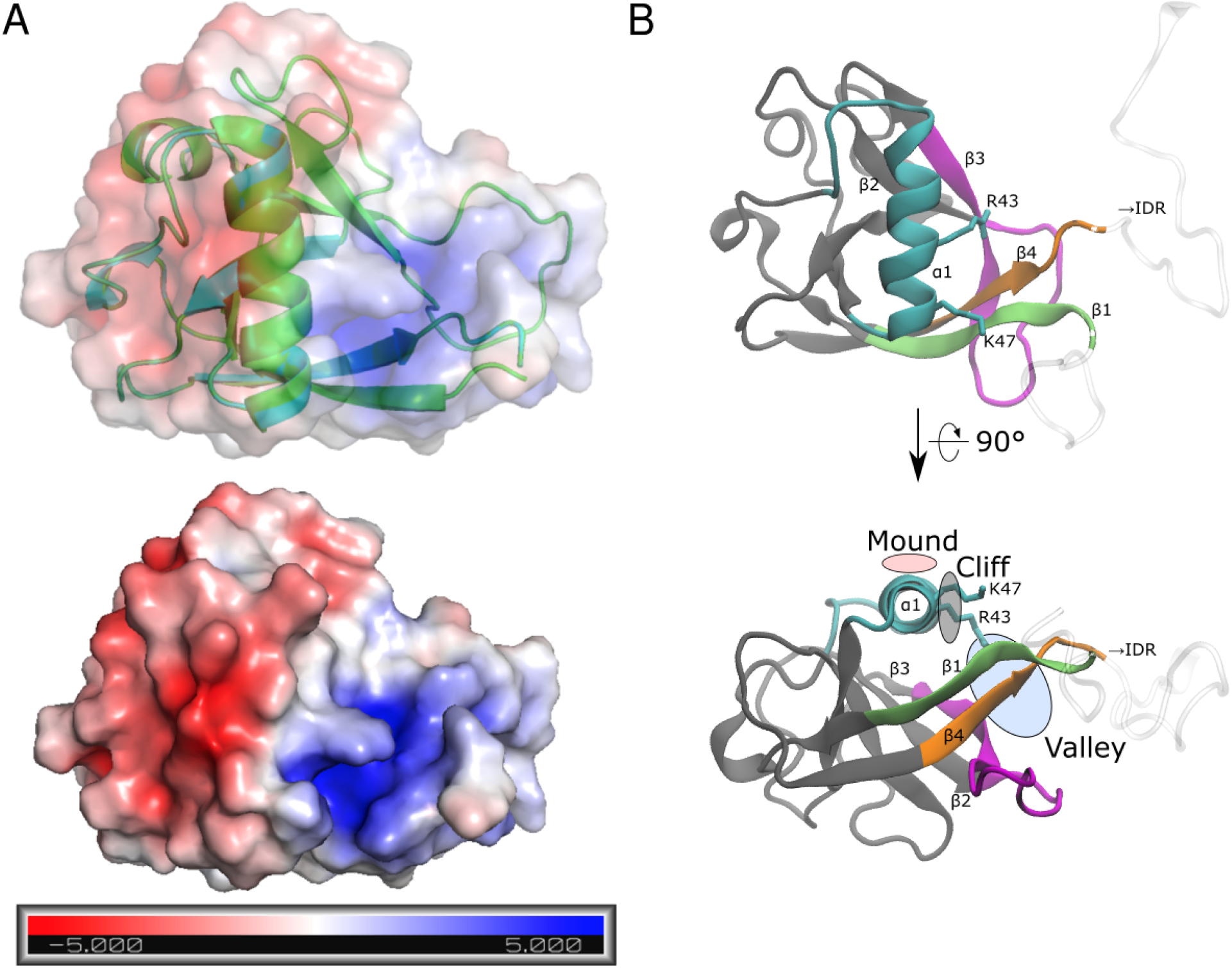
(A) Surface electrostatic potential of the nsp1 and (B) annotated surface structure of the nsp1 recognition sites for SL1. In (A), units are in *k_B_T/e*, where *k_B_* is the Boltzmann factor, *T* is the temperature of the system (= 300 K), and *e* is the unit charge of a proton. Color coding in (B) corresponds to the region defined in Fig. 1 D.

Although the binding site for SL1 on nsp1 can be characterized as an interface consisting of regions (i) through (v), SL1 did not assume a stable conformation, even when it was bound to these regions. Diverse binding modes were observed in the canonical ensemble. Although SL1 nearly always interacted with residues in the region (iv), its conformation was diverse and fluctuated greatly. In addition, the nsp1 IDR was also highly flexible.

### Clustering analysis of the binding poses

The diversity of the binding modes was further investigated using cluster analysis based on the contact map for each snapshot (see the Methods section). We determined the clustering threshold using the criterion that any cluster has at least one inter-residue contacts with more than 80 % in each cluster. As a result, the binding modes could be categorized into 14 clusters and outliers, which had 34.2% of the statistical weight in the canonical ensemble. In even the most major cluster, the statistical weight was only 15.5%; those for the second, third, and fourth clusters were 9.9%, 7.4%, and 5.0%, respectively. Each cluster had a unique tendency to use a set of binding regions (Supplementary Table S2, Supplementary Figure S3). We also analyzed the differences in surface areas of the interacting interfaces in the ordered and disordered regions of nsp1 among the 14 clusters (Supplementary Figure S4). The distribution shows the unique characteristics of each cluster. These results indicate that recognition of SL1 by nsp1 is established by multimodal binding modes.

The representative structure of cluster 1, which had the largest population among all clusters, is presented in Fig. 7 and Supplementary Table S3. Nsp1 recognized SL1 via regions (ii), (iii) and (iv). In the region (ii), the basic residues in H2 formed the Arg43–C17 and Lys47–U16 salt-bridges. Region (iii) recognized SL1 via the Asp75–U18 hydrogen bond. Residues Arg124 through Gly137 in region (iv) attached to SL1 via the Arg124–U17, Ala131–C19, and Ser135–C16 hydrogen bonds; Tyr136 stacks between C21 and G23 instead of A22, which was flipped out. Representative structures of clusters 2 and 3 are also presented in the supporting material (Supplementary Figures S5 and S6 and Supplementary Tables S4 and S5).

**Figure 7:**
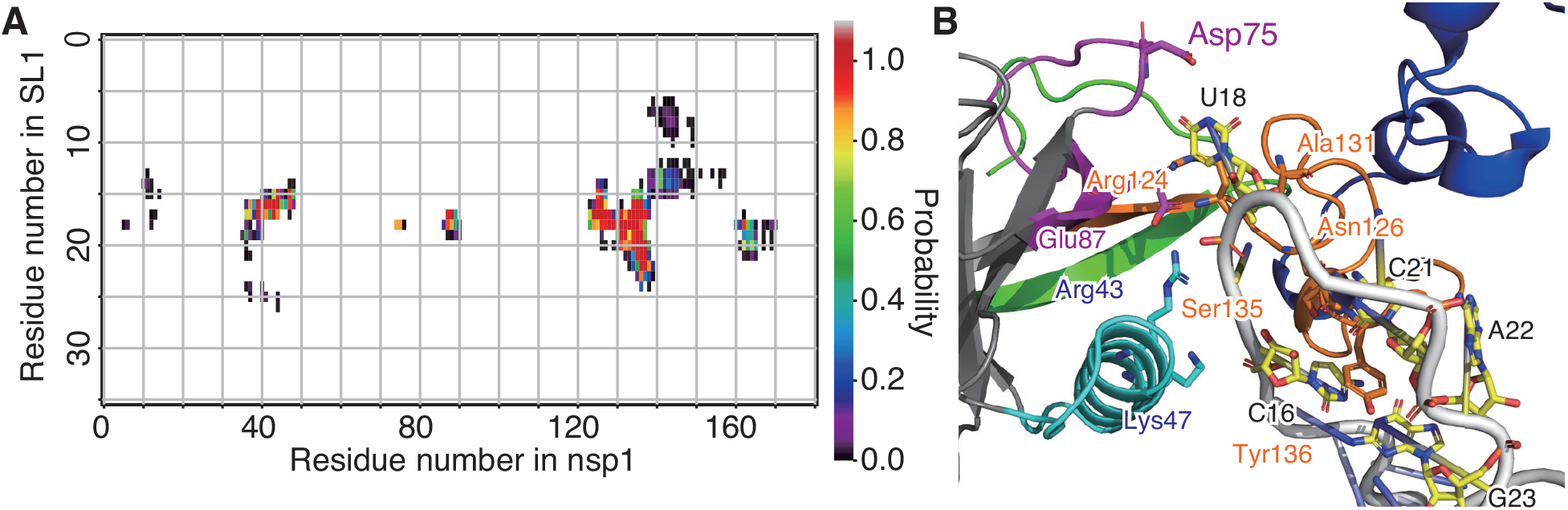
Interactions between nsp1 and SL1 observed in cluster 1. (A) Pairwise contact probability in cluster 1. See the legend to Fig. 4. (B) Representative snapshot of the cluster 1. The interface regions (i) through (v) are shown as green, cyan, magenta, red, and blue ribbons. Bases 16–26 of SL1 are shown in orange.

### Model of the mechanism

Based on the simulation results, we suggest a model of the mechanism in Fig. 8. Without SL1, both the N-terminal domain and C-terminal helices bind to the 40*S* ribosome, blocking human mRNAs. With the SL1, the binding between SL1 and nsp1 at both the N-terminal domain and part of the IDR results in the C-terminal helices being pulled such that they can no longer maintain their binding to the 40*S* ribosome. The 5’-end of the viral RNA will then be loaded into the ribosome, initiating the translation.

**Figure 8:**
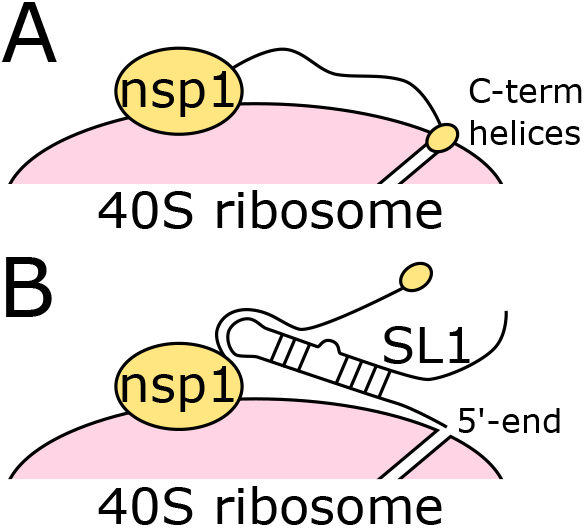
Schematic model of the nsp1 translation shutoff evasion suggested by our simulation. (A) Without SL1, the C-terminal helices of nsp1 obstruct the tunnel for mRNA. (B) With SL1, the N-terminal domain and the nsp1 IDR bind to SL1, and the C-terminal helices dissociate from the ribosome, opening the tunnel to mRNA.

### Relation to other experimental results

It has been reported that the Arg124Ala– Lys125Ala double nsp1 mutant lacks the ability to recognize viral RNA.^3^ This can be explained by the results of our simulation, which showed that sidechain of Arg124 strongly interacts with the phosphate backbone of U18 (Table 1 and Figs. 5 and 7). An Arg124Ala mutation would eliminate the ionic interaction between the sidechain and the backbone, and nsp1 would lose its ability to recognize viral RNA.

The circular dichroism spectrum of the SARS-CoV-2 nsp1 C-terminal region (residues 130-180)^16^ in solution had only a single peak at 198 nm and did not show ellipticities at 208 nm and 222 nm. This indicates that the nsp1 C-terminal region did not form *α*-helices or *β*-sheets and was disordered. Although in our simulation we found that nsp1 partially forms the *α*-helix in the IDR, our simulation also indicated that the percentage of the helix formed was low and the structure was unstable, which may explain the difference from the experimental results. The difference may also be attributable to the presence of RNA and other solvent conditions and the force field inaccuracies. Further study will be needed before a conclusion can be drawn.

Whether SARS-CoV-2 nsp1 and SL1 bind without the ribosome is controversial. It has been reported that nsp1 and bases 7-33 of SARS-CoV-2 bind with a binding constant of 0.18 μM,^49^ but it has also been reported that a gel shift does not occur with the 5’-UTR of SARS-CoV-2 at concentrations up to 20 μM when tRNAs was used to exclude the non-specific binding.^6^ The present simulation results indicate that the binding mode observed herein did not have a specific, defined structure. Typically, with such binding modes, the binding is expected to be weak; we therefore do not think these simulation results contradict the results from either of the aforementioned experiments.

Mutations to SL1 bases 14-25, which disrupt the Watson-Crick pairs of the stem loop, reportedly cause translation to be shut off.^6^ That observation is consistent with our finding that the hairpin structure of bases 18-22 in SL1 is recognized by nsp1. Hydrogen-bond interaction analysis showed that the RNA phosphate backbone is mainly recognized within the C15-C20 region (Table 1 and Fig. 5). Moreover, our finding is consistent with the fact that the sequence of the hairpin region (corresponding to U18-C21 in our simulation) is not well conserved among SARS-CoV-2 mutational variants, whereas that of the stem is well conserved. ^50^ Our simulation shows that the interaction between nsp1 and the SL1 backbone is stronger than that between nsp1 and the SL1 sidechains (Table 1), which highlights the importance of the backbone interaction.

### Limitations of this study

Our simulation was performed based on several assumptions. Here, we list the limitations of the present study.

First, our simulation was performed without the ribosome. This was mainly because the simulation started before the structure of the nsp1-ribosome complex was deposited. Furthermore, the orientation of the nsp1 N-terminal domain attached to the 40*S* ribosome is still ambiguous in density maps. With the 40*S* ribosome, the environment around nsp1 may be altered and so be the interaction between the RNA and nsp1. Specifically, the ribosome consists mostly of ribosomal RNAs and are therefore strongly negatively charged, which may change the interaction environment significantly.

To maintain the stability of the hairpin loop structure, we performed the simulation with restraints on the G-C pairs in the 5’-UTR. These restraints may have hindered RNA forming structures other than the initial hairpin structure. However, in the secondary structure prediction using CentroidFold,^51^ these base pairs were predicted to exist in more than 92% of the ensemble. Furthermore, a recent study^52^ showed that, even with a rigorous extended ensemble simulation, the hairpin structure remained intact. Given these results, the drawback of structural restraints to SL1 is expected to be minimal.

Finally, as is always the case with a simulation study, the mismatch between the simulation force field and the real world leaves a non-negligible gap. In addition, some residues may have alternative protonation states upon binding to RNA (e.g., histidine protonation state).

### Future research and conclusions

The present simulation was performed with only nsp1 and SL1. Arguably, simulation of a complex consisting of the 40*S* ribosome, nsp1 and SL1 will be an important step toward further understanding the details of the mechanism underlying the evasion of nsp1 by viral RNA. Our results show that nsp1-SL1 binding has multimodal binding structures. The addition of the 40*S* ribosome to the system may restrict the structure to a smaller number of possible binding poses and possibly tighter binding poses may be obtained.

In addition to a simulation study, mutational analysis of nsp1 will be informative. In addition to the already known mutation at Arg124, current simulation results predict Lys47, Arg43, and Asn126 are important to nsp1-SL1 bninding. Mutation analyses of these residues will help us to understand the molecular mechanism of nsp1.

Finally, the development of inhibitors of nsp1-stem loop binding, is highly anticipated in the current pandemic. Although the present results imply that a specific binding structure might not exist, important residues in nsp1 and bases in SL1 were detected. Blocking or mimicking the binding of these residues/bases, could potentially nullify the function of nsp1.

In conclusion, using MD simulation, we investigated the binding and molecular mechanism of SARS-CoV-2 nsp1 and the 5’-UTR stem loop of SARS-CoV-2. The results suggest that the interaction between nsp1 and the 5’-UTR stem loop prevents the C-terminal helices from binding to the ribosome, thereby avoiding translation shutoff. The interaction analysis further revealed that the hairpin loop structure of the 5’-UTR stem loop is recognized by both the N-terminal domain and the intrinsically disordered region of nsp1. Multiple binding poses were obtained, and the largest cluster of binding poses included interactions that coincide with the results of the previous mutational analysis.

## Supporting information

Additional force field file used in the simulation

Additional modifications applied to the simulation program (GROMACS 2016)

## Acknowledgement

We thank Dr. Atsushi Matsumoto for his technical assistance. This research was supported by the Basis for Supporting Innovative Drug Discovery and Life Science (BINDS) project under Grant Number JP20am0101106j0004, Agency for Medical Research and Development (AMED), Japan to HK; by a Grant-in-Aid for Early-Career Scientists from the Japan Society for the Promotion of Science (JSPS), Japan to SS (JP16K17778), by a Grant-in-Aid for Scientific Research (A) from the JSPS to SS (JP16H02484), (C) to KK (JP20K12069), and (C) to JI (JP20K12041), and by a Grant-in-Aid for Scientific Research on Innovative Areas from the Ministry of Education, Culture, Sports, Science and Technology (MEXT) to SS (JP19H05410). Simulations were performed on supercomputers at Research Center for Computational Science, Okazaki, and Academic Center for Computing and Media Studies, Kyoto University. Authors declare no competing interests.

## Supporting Information Available

The supporting information is available free of charge:

- Assessment of the convergence, detailed procedure of the center-of-mass distance from the Cryo-EM density map, and detailed description of clusters. (SI.pdf)
- Parameter files for the RNA-cap force field (m7G-ppp-(2’-O-Me-A)). (forcefield.zip)
- Modifications added to the software as a series of patch files. (patch.zip)

## Notes

https://bsma.pdbj.org/entry/26

### Competing Interest Statement

The authors have declared no competing interest.

### Summary of Updates

Overall quality improvements

## References

(1) Narayanan, K.; Huang, C.; Lokugamage, K.; Kamitani, W.; Ikegami, T.; Tseng, C.-T. K.; Makino, S. Severe Acute Respiratory Syndrome Coronavirus nsp1 Suppresses Host Gene Expression, Including That of Type I Interferon, in Infected Cells. Journal of Virology 2008, 82, 4471–4479.

(2) Kamitani, W.; Huang, C.; Narayanan, K.; Lokugamage, K. G.; Makino, S. A two-pronged strategy to suppress host protein synthesis by SARS coronavirus Nsp1 protein. Nature Structural & Molecular Biology 2009, 16, 1134–1140.

(3) Lokugamage, K. G.; Narayanan, K.; Huang, C.; Makino, S. Severe Acute Respiratory Syndrome Coronavirus Protein nsp1 Is a Novel Eukaryotic Translation Inhibitor That Represses Multiple Steps of Translation Initiation. Journal of Virology 2012, 86, 13598–13608.

(4) Tanaka, T.; Kamitani, W.; DeDiego, M. L.; Enjuanes, L.; Matsuura, Y. Severe Acute Respiratory Syndrome Coronavirus nsp1 Facilitates Efficient Propagation in Cells through a Specific Translational Shutoff of Host mRNA. Journal of Virology 2012, 86, 11128–11137.

(5) Narayanan, K.; Ramirez, S. I.; Lokugamage, K. G.; Makino, S. Coronavirus nonstructural protein 1: Common and distinct functions in the regulation of host and viral gene expression. Virus Research 2015, 202, 89–100.

(6) Tidu, A.; Janvier, A.; Schaeffer, L.; Sosnowski, P.; Kuhn, L.; Hammann, P.; Westhof, E.; Eriani, G.; Martin, F. The viral protein NSP1 acts as a ribosome gatekeeper for shutting down host translation and fostering SARS-CoV-2 translation. RNA 2020, in press.

(7) Kamitani, W.; Narayanan, K.; Huang, C.; Lokugamage, K.; Ikegami, T.; Ito, N.; Kubo, H.; Makino, S. Severe acute respiratory syndrome coronavirus nsp1 protein suppresses host gene expression by promoting host mRNA degradation. Proceedings of the National Academy of Sciences 2006, 103, 12885–12890.

(8) Huang, C.; Lokugamage, K. G.; Rozovics, J. M.; Narayanan, K.; Semler, B. L.; Makino, S. SARS Coronavirus nsp1 Protein Induces Template-Dependent Endonucleolytic Cleavage of mRNAs: Viral mRNAs Are Resistant to nsp1-Induced RNA Cleavage. PLoS Pathogens 2011, 7, e1002433.

(9) Wathelet, M. G.; Orr, M.; Frieman, M. B.; Baric, R. S. Severe Acute Respiratory Syndrome Coronavirus Evades Antiviral Signaling: Role of nsp1 and Rational Design of an Attenuated Strain. Journal of Virology 2007, 81, 11620–11633.

(10) Thoms, M. et al. Structural basis for translational shutdown and immune evasion by the Nsp1 protein of SARS-CoV-2. Science 2020, 369, 1249–1255.

(11) Schubert, K.; Karousis, E. D.; Jomaa, A.; Scaiola, A.; Echeverria, B.; Gurzeler, L.-A.; Leibundgut, M.; Thiel, V.; Mühlemann, O.; Ban, N. SARS-CoV-2 Nsp1 binds the ribosomal mRNA channel to inhibit translation. Nature Structural & Molecular Biology 2020, 27, 959–966.

(12) Yuan, S.; Peng, L.; Park, J. J.; Hu, Y.; Devarkar, S. C.; Dong, M. B.; Shen, Q.; Wu, S.; Chen, S.; Lomakin, I. B.; Xiong, Y. Nonstructural Protein 1 of SARS-CoV-2 Is a Potent Pathogenicity Factor Redirecting Host Protein Synthesis Machinery toward Viral RNA. Molecular Cell 2020, 80, 1055–1066.e6.

(13) Kim, D.; Lee, J.-Y.; Yang, J.-S.; Kim, J. W.; Kim, V. N.; Chang, H. The Architecture of SARS-CoV-2 Transcriptome. Cell 2020, 181, 914–921.e10.

(14) Banerjee, A. K. et al. SARS-CoV-2 Disrupts Splicing, Translation, and Protein Trafficking to Suppress Host Defenses. Cell 2020, 183, 1325–1339.e21.

(15) Almeida, M. S.; Johnson, M. A.; Herrmann, T.; Geralt, M.; Wüthrich, K. Novel *β*-Barrel Fold in the Nuclear Magnetic Resonance Structure of the Replicase Nonstructural Protein 1 from the Severe Acute Respiratory Syndrome Coronavirus. Journal of Virology 2007, 81, 3151–3161.

(16) Kumar, A.; Kumar, A.; Kumar, P.; Garg, N.; Giri, R. SARS-CoV-2 NSP1 C-terminal region (residues 130-180) is an intrinsically disordered region. bioRxiv 2020,

(17) Fiser, A.; Šali, A. Methods in Enzymology; Elsevier, 2003; pp 461–491.

(18) Cornell, W. D.; Cieplak, P.; Bayly, C. I.; Gould, I. R.; Merz, K. M.; Ferguson, D. M.; Spellmeyer, D. C.; Fox, T.; Caldwell, J. W.; Kollman, P. A. A second generation force field for the simulation of proteins, nucleic acids, and organic molecules. Journal of the American Chemical Society 1995, 117, 5179–5197.

(19) Wang, J.; Cieplak, P.; Kollman, P. A. How well does a restrained electrostatic potential (RESP) model perform in calculating conformational energies of organic and biological molecules? Journal of Computational Chemistry 2000, 21, 1049–1074.

(20) Hornak, V.; Abel, R.; Okur, A.; Strockbine, B.; Roitberg, A.; Simmerling, C. Comparison of multiple Amber force fields and development of improved protein backbone parameters. Proteins: Structure, Function, and Bioinformatics 2006, 65, 712–725.

(21) Maier, J. A.; Martinez, C.; Kasavajhala, K.; Wickstrom, L.; Hauser, K. E.; Simmerling, C. ff14SB: Improving the Accuracy of Protein Side Chain and Backbone Parameters from ff99SB. Journal of Chemical Theory and Computation 2015, 11, 3696–3713.

(22) Popenda, M.; Szachniuk, M.; Antczak, M.; Purzycka, K. J.; Lukasiak, P.; Bartol, N.; Blazewicz, J.; Adamiak, R. W. Automated 3D structure composition for large RNAs. Nucleic Acids Research 2012, 40, e112–e112.

(23) Antczak, M.; Popenda, M.; Zok, T.; Sarzynska, J.; Ratajczak, T.; Tomczyk, K.; Adamiak, R. W.; Szachniuk, M. New functionality of RNAComposer: application to shape the axis of miR160 precursor structure. Acta Biochimica Polonica 2017, 63.

(24) Wu, F. et al. A new coronavirus associated with human respiratory disease in China. Nature 2020, 579, 265–269.

(25) Bayly, C. I.; Cieplak, P.; Cornell, W.; Kollman, P. A. A well-behaved electrostatic potential based method using charge restraints for deriving atomic charges: the RESP model. J. Phys. Chem. 1993, 97, 10269–10280.

(26) Perez, A.; Marchan, I.; Svozil, D.; Sponer, J.; Cheatham, T. E.; Laughton, C. A.; Orozco, M. Refinement of the AMBER force field for nucleic acids: improving the description of alpha/gamma conformers. Biophys. J. 2007, 92, 3817–3829.

(27) Zgarbová, M.; Otyepka, M.; Šponer, J.; Mladek, A.; Banas, P.; Cheatham, T. E.; Jurecka, P. Refinement of the Cornell et al. Nucleic Acids Force Field Based on Reference Quantum Chemical Calculations of Glycosidic Torsion Profiles. J. Chem. Theory Comput. 2011, 7, 2886–2902.

(28) da Silva, A. W. S.; Vranken, W. F. ACPYPE - AnteChamber PYthon Parser interfacE. BMC Research Notes 2012, 5, 367.

(29) Case, D. et al. AMBER 2017. University of California, San Francisco, 2017.

(30) Jorgensen, W. L.; Chandrasekhar, J.; Madura, J. D.; Impey, R. W.; Klein, M. L. Comparison of simple potential functions for simulating liquid water. The Journal of Chemical Physics 1983, 79, 926–935.

(31) Joung, I. S.; Cheatham, T. E. Determination of alkali and halide monovalent ion parameters for use in explicitly solvated biomolecular simulations. J. Phys. Chem. B 2008, 112, 9020–9041.

(32) Ikebe, J.; Sakuraba, S.; Kono, H. H3 histone tail conformation within the nucleosome and the impact of K14 acetylation studied using enhanced sampling simulation. PLoS computational biology 2016, 12, e1004788.

(33) Li, Z.; Kono, H. Investigating the Influence of Arginine Dimethylation on Nucleosome Dynamics Using All-Atom Simulations and Kinetic Analysis. The Journal of Physical Chemistry B 2018, 122, 9625–9634.

(34) Kasahara, K.; Shiina, M.; Higo, J.; Ogata, K.; Nakamura, H. Phosphorylation of an intrinsically disordered region of Ets1 shifts a multi-modal interaction ensemble to an auto-inhibitory state. Nucleic acids research 2018, 46, 2243–2251.

(35) Wang, L.; Friesner, R. A.; Berne, B. J. Replica Exchange with Solute Scaling: A More Efficient Version of Replica Exchange with Solute Tempering (REST2). The Journal of Physical Chemistry B 2011, 115, 9431–9438.

(36) Abraham, M. J.; Murtola, T.; Schulz, R.; Páll, S.; Smith, J. C.; Hess, B.; Lindahl, E. GROMACS: High performance molecular simulations through multi-level parallelism from laptops to supercomputers. SoftwareX 2015, 1, 19–25.

(37) Bussi, G. Hamiltonian replica exchange in GROMACS: a flexible implementation. Molecular Physics 2013, 112, 379–384.

(38) Bussi, G.; Donadio, D.; Parrinello, M. Canonical sampling through velocity rescaling. The Journal of Chemical Physics 2007, 126, 014101.

(39) Hess, B.; Bekker, H.; Berendsen, H. J. C.; Fraaije, J. G. E. M. LINCS: A linear constraint solver for molecular simulations. Journal of Computational Chemistry 1997, 18, 1463–1472.

(40) Souaille, M.; Roux, B. Extension to the weighted histogram analysis method: combining umbrella sampling with free energy calculations. Computer Physics Communications 2001, 135, 40–57.

(41) Shirts, M. R.; Chodera, J. D. Statistically optimal analysis of samples from multiple equilibrium states. J. Chem. Phys. 2008, 129, 124105.

(42) Humphrey, W.; Dalke, A.; Schulten, K. VMD – Visual Molecular Dynamics. Journal of Molecular Graphics 1996, 14, 33–38.

(43) Schrödinger, LLC,

(44) Kabsch, W.; Sander, C. Dictionary of protein secondary structure: Pattern recognition of hydrogen-bonded and geometrical features. Biopolymers 1983, 22, 2577–2637.

(45) McGibbon, R. T.; Beauchamp, K. A.; Harrigan, M. P.; Klein, C.; Swails, J. M.; Hernández, C. X.; Schwantes, C. R.; Wang, L.-P.; Lane, T. J.; Pande, V. S. MD-Traj: A Modern Open Library for the Analysis of Molecular Dynamics Trajectories. Biophysical Journal 2015, 109, 1528–1532.

(46) Ester, M.; Kriegel, H.-P.; Sander, J.; Xu, X., et al. A density-based algorithm for discovering clusters in large spatial databases with noise. KDD. 1996; pp 226–231.

(47) Ankerst, M.; Breunig, M. M.; Kriegel, H.-P.; Sander, J. OPTICS. ACM SIGMOD Record 1999, 28, 49–60.

(48) Semper, C.; Watanabe, N.; Savchenko, A. Structural characterization of nonstructural protein 1 from SARS-CoV-2. iScience 2021, 24, 101903.

(49) Vankadari, N.; Jeyasankar, N. N.; Lopes, W. J. Structure of the SARS-CoV-2 Nsp1/5′-Untranslated Region Complex and Implications for Potential Therapeutic Targets, a Vaccine, and Virulence. The Journal of Physical Chemistry Letters 2020, 11, 9659–9668.

(50) Miao, Z.; Tidu, A.; Eriani, G.; Martin, F. Secondary structure of the SARS-CoV-2 5’-UTR. RNA Biology 2020, 1–10.

(51) Sato, K.; Hamada, M.; Asai, K.; Mituyama, T. CENTROIDFOLD: a web server for RNA secondary structure prediction. Nucleic Acids Research 2009, 37, W277–W280.

(52) Bottaro, S.; Bussi, G.; Lindorff-Larsen, K. Conformational Ensembles of Non-Coding Elements in the SARS-CoV-2 Genome from Molecular Dynamics Simulations. bioRxiv 2020,.

