## Supporting Information for "Modeling the SARS-CoV-2 nsp1–5’-UTR complex via extended ensemble simulations"

#### Convergence of the simulations

We monitored the convergence of the simulation by two measures: the secondary structure and the hydrogen bond forming rate. Figure S1 shows the probability distribution of the nsp1 secondary structure among different simulation lengths. We plotted the secondary structure forming probability using trajectories at 5–10 ns, 10–20 ns, 15–30 ns, 20–40 ns, and 25–50 ns of the simulation in replica 0. The secondary structure converged after 15–30 ns. We also tested the hydrogen bond forming rate in Fig. S2. We took trajectories from $x$ ns to $2x$ ns, e.g. 15 ns to 30 ns for the case of $x = 15$, and the probability of the hydrogen bonds between nsp1 and SL1 was assessed. Similar to the secondary structure, the hydrogen bond forming rates converged after $x \approx 15$ ns. From these results, we used 25–50 ns of the simulation in the subsequent analyses.

### Modeling of nsp1-40S ribosome structure

To further investigate the structure of the nsp1, and to investigate the geometrical restriction in an nsp1-40S ribosome complex, we modeled an nsp1-40S ribosome complex by the density fitting approach. We fitted the SARS-CoV-2 nsp1 N-terminal domain structure (PDB ID: 7K3N) into the ribosome–nsp1 complex cryo-EM density map (Electron Microscopy Data Bank ID: EMD-11276) using the structure of 40S ribosome–nsp1 C-terminal helices complex (PDB ID: 6ZLW). We used UCSF Chimera[1] to fit the density map. Six models over the correlation coefficient of 0.80 were selected for further analysis. After the fitting, we measured the distance between the center-of-mass of the globular domain (residues 14 to 125) C$\alpha$ atoms and the center-of-mass of the C-terminal helices domain (residues 153 to 179) C$\alpha$ atoms, and averaged the distances among 6 models.

### Characterstics of clusters

Table S1 lists the residues involved in nsp1–SL1 binding. Figure S3 shows the structure of all 14 clusters identified in our analysis. Detailed characteristics of the clusters are listed in Table S2 and Fig. S4.

### Details of clusters 2 and 3

Cluster 2 interacted with SL1 via the interface regions (i), (ii), and (iv) (Fig. S5. A remarkable feature of cluster 2 is the recognition of C19, C20, and C21. Arg43 and Lys47 in the region (ii) formed the salt-bridge with their backbone in 70.6 % and 100.0 % in cluster 2. The region (iv). The base of C21 was fliped out from the stem loop was stacked Asn126 amide group and Gly137 backbone. Asn126 also formed hydrogen bonds with C19 (96.7 %) and C20 (92.3 %).

Cluster 3 showed interactions with a wider range of SL1 than others. Lys11 and Lys125 frequently formed salt-bridges with U13 (91.5 %) and A14 (88.8 %), respectively. His134 showed a high contact frequency with C32. In addition, the side chain of Asp126 entered the center of the stem loop and formed the hydrogen bonds with the bases of U17 and C20.

### References

(1) Pettersen, E. F.; Goddard, T. D.; Huang, C. C.; Couch, G. S.; Greenblatt, D. M.; Meng, E. C.; Ferrin, T. E. UCSF Chimera–a visualization system for exploratory research and analysis. *Journal of computational chemistry* **2004**, *25*, 1605–1612.

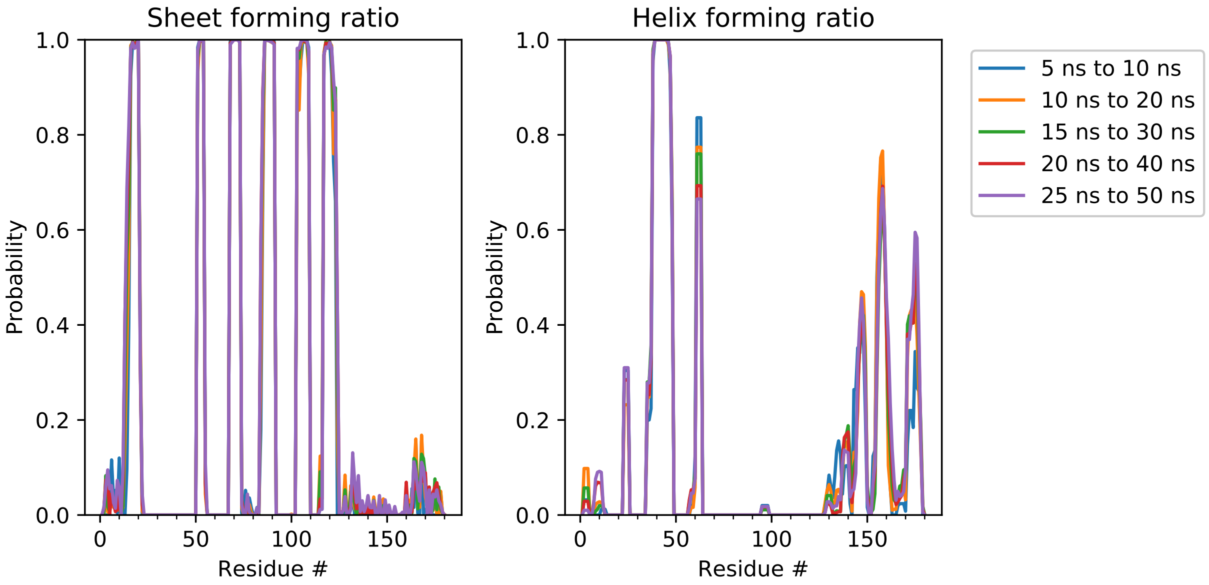

Figure S1: Secondary structure distribution of the simulation corresponding to replica number 0. "Helix" includes the DSSP assignment of $\alpha$-helix, 3-10 helix and $\pi$-helix. "Sheet" includes both $\beta$-bridge and $\beta$-ladder.

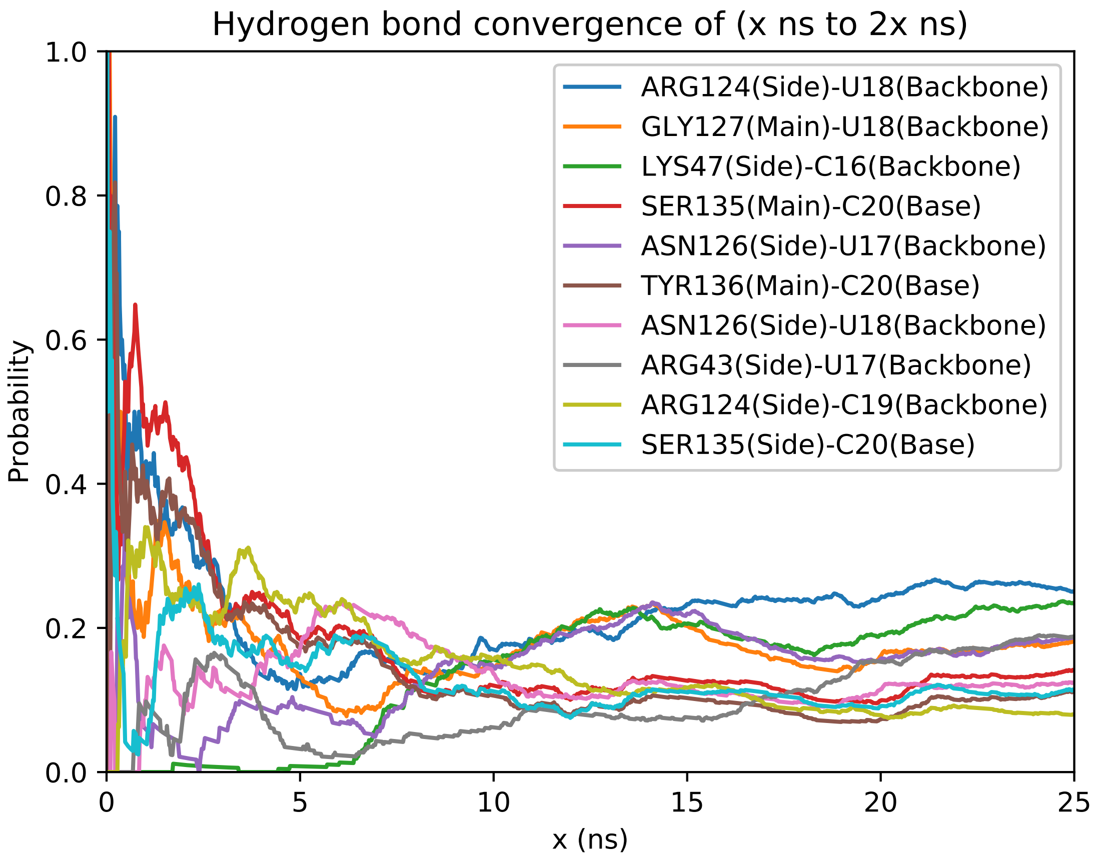

Figure S2: Hydrogen bond forming rate between $x$ ns and $2x$ ns of the simulation in replica 0. Hydrogen bonds having the 10 largest probabilities among 0–50 ns simulation were selected and plotted in the figure.

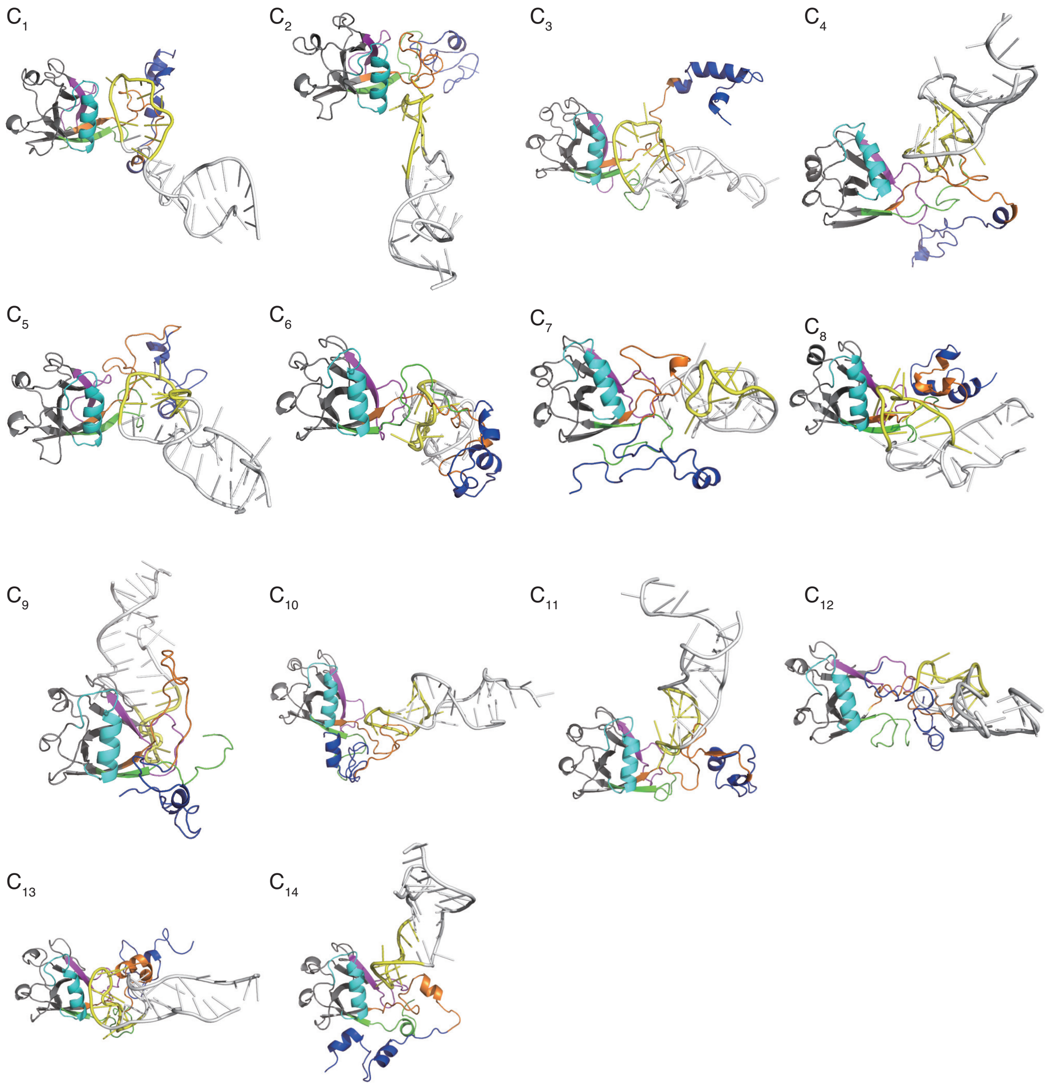

Figure S3: Examples of a snapshot in each cluster. The panels ($C_1$) through ($C_{14}$) show snapshots of clusters 1 through 14, respectively. The interface regions (i) through (v) are shown in green, cyan, magenta, orange, and blue ribbons. The 16–26th bases of SL1 are shown in yellow.

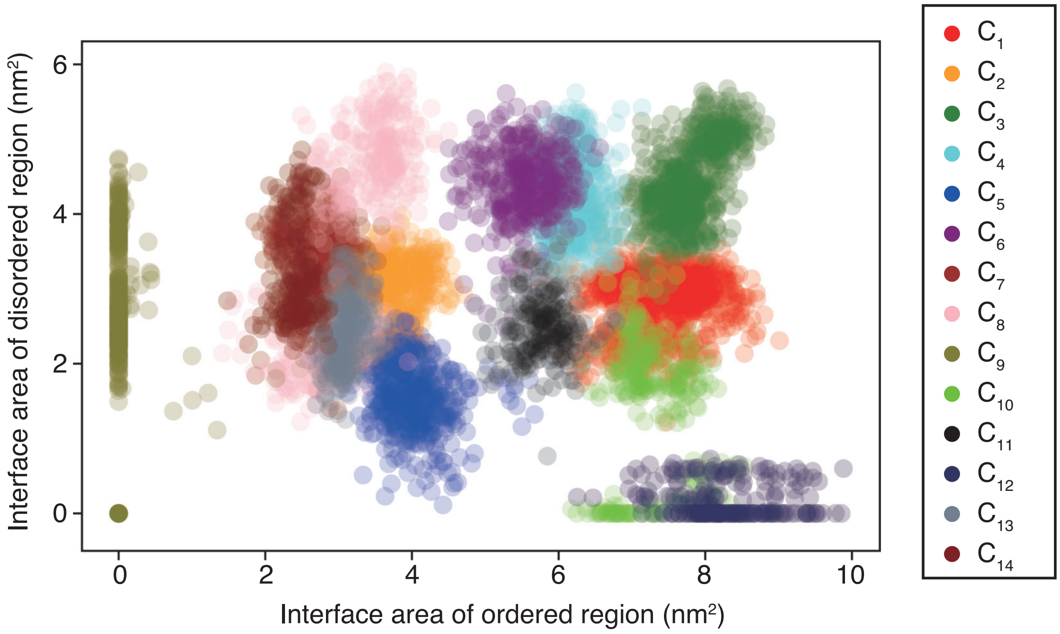

Figure S4: The surface area of interaction interfaces of Nsp1 in each snapshot. $C_1$ through $C_{14}$ represent clusters numbered 1 to 14, respectively. The horizontal and vertical axes indicate the area on ordered and disordered regions, respectively. Color corresponds to the cluster ID.

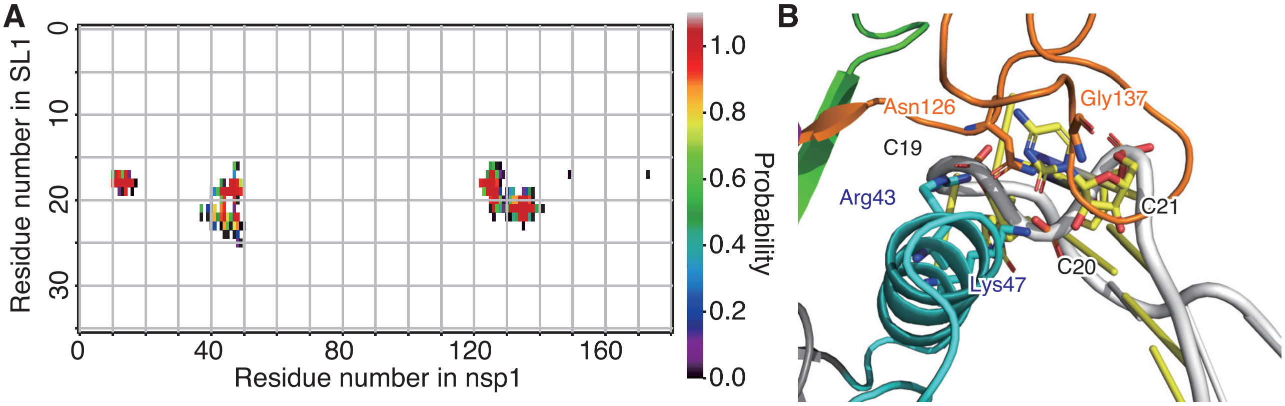

Figure S5: Interactions between Nsp1 and SL1 in cluster 2. (A) Pairwise contact probability for each cluster. See the legend of Fig. 4 in the main paper. (B) A snapshot in cluster 2.

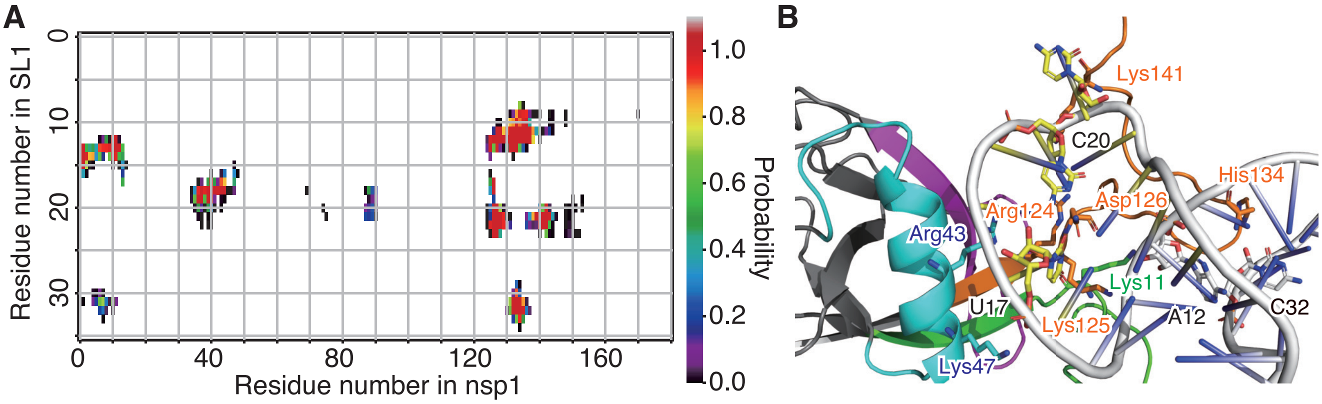

Figure S6: Interactions between Nsp1 and SL1 in cluster 3. (A) Pairwise contact probability for each cluster. See the legend of Fig. 4 in the main paper. (B) A snapshot in cluster 3.

Table S1: Characterization of SL1 binding regions of Nsp1.

| Region ID | Residues | %* |
|---|---|---|
| (i) | 1–18 | 72.1 |
| (ii) | 31–50 | 81.8 |
| (iii) | 74–90 | 63.2 |
| (iv) | 121–146 | 97.4 |
| (v) | 147–180 | 59.2 |

* The probability to form at least one contact between the region and SL1.

Table S2: Characteristics of each conformational cluster.

| Cluster ID | Total% | (i)% | (ii)% | (iii)% | (iv)% | (v)% | Representative contacts |
|---|---|---|---|---|---|---|---|
| 1 | 15.5 | 14.0 | 100 | 100 | 100 | 98.5 | Arg124–U17 |
| 2 | 9.88 | 100 | 100 | 0 | 100 | 1.77 | Lys47–C20 |
| 3 | 7.42 | 100 | 100 | 81.4 | 100 | 6.96 | Arg126–C20 |
| 4 | 4.98 | 100 | 100 | 100 | 100 | 0 | His134–C20 |
| 5 | 4.91 | 37.5 | 100 | 91.0 | 100 | 100 | Gly127–U18 |
| 6 | 4.88 | 100 | 0 | 100 | 100 | 80.6 | Arg77–C20 |
| 7 | 3.98 | 100 | 0 | 100 | 100 | 0.91 | Ser142–A22 |
| 8 | 3.60 | 100 | 100 | 24.3 | 100 | 8.16 | Arg43–C19 |
| 9 | 2.62 | 13.0 | 0.88 | 87.8 | 0.88 | 0 | Arg73–A22 |
| 10 | 2.30 | 52.7 | 77.5 | 86.0 | 100 | 100 | Asn162-U18 |
| 11 | 1.85 | 100 | 100 | 100 | 100 | 13.8 | Ser135–C20 |
| 12 | 1.36 | 0 | 0 | 34.0 | 100 | 100 | Ser135–A22 |
| 13 | 1.31 | 75.5 | 100 | 45.7 | 100 | 0 | Gly127–C19 |
| 14 | 1.20 | 0 | 100 | 100 | 100 | 1.74 | Arg43–C20 |
| outliers | 34.2 | 84.6 | 85.7 | 41.0 | 100 | 88.1 | - |

Table S3: Hydrogen bonds and salt-bridges in cluster 1.

| Detected interactions | % | Bond type |
|---|---|---|
| Arg124–U17 | 96.0 | H-bond |
| Asp75–U18 | 95.2 | H-bond |
| Ser40–U17 | 84.3 | H-bond |
| Ala131–C19 | 78.8 | H-bond |
| Ser135–C16 | 78.4 | H-bond |
| Arg124–U18 | 72.0 | H-bond |
| Lys47–C16 | 71.2 | H-bond |
| Asn126–C16 | 59.0 | H-bond |
| Arg43–U17 | 56.4 | H-bond |
| Lys47–C16 | 82.3 | salt-bridge |
| Arg43–U17 | 81.1 | salt-bridge |
| Arg43–U18 | 76.2 | salt-bridge |
| Arg124–U17 | 75.2 | salt-bridge |
| Lys141–A14 | 57.4 | salt-bridge |

Table S4: Hydrogen bonds and salt-bridges in cluster 2.

| Detected bonds | % | Bond type |
|---|---|---|
| Lys47–C20 | 96.9 | hbond |
| Asn126–C19 | 96.7 | hbond |
| Asn126–C20 | 92.3 | hbond |
| Lys47–U18 | 82.3 | hbond |
| Arg43–C20 | 69.6 | hbond |
| Ser40–C21 | 63.8 | hbond |
| Gln44–C20 | 55.5 | hbond |
| Lys047–C20 | 100.0 | salt-bridge |
| Arg043–C20 | 70.6 | salt-bridge |

Table S5: Hydrogen bonds and salt-bridges in cluster 3.

| Detected bonds | % | Bond type |
| --- | --- | --- |
| Asn126–C20 | 96.5 | hbond |
| Asn126–U17 | 87.0 | hbond |
| Lys141–C21 | 85.8 | hbond |
| Lys125–A14 | 77.3 | hbond |
| Lys11–U13 | 77.1 | hbond |
| Ser142–A22 | 72.8 | hbond |
| Met1–C15 | 70.7 | hbond |
| Ser135–U11 | 65.2 | hbond |
| Ser40–U18 | 56.2 | hbond |
| Asn126–G23 | 55.6 | hbond |
| Lys011–U13 | 91.5 | salt-bridge |
| Lys125–A14 | 88.8 | salt-bridge |
