## Additional modifications applied to the simulation program (GROMACS 2016) for "Modeling the SARS-CoV-2 nsp1–5’-UTR complex via extended ensemble simulations": readme.txt

Usage: 
1) Apply patch to gromacs 2016 by the patch command:
patch -p1 < patch_gromacs_2016.patch

2) Compile and install GROMACS as usual. For numerical precision problem I recommend double precision.

3) Run gromacs with MPI and -hrex option, e.g.
mpirun -np 64 gmx_d_mpi mdrun -deffnm rest -multidir repl0 repl1 repl2 repl3 -replex 1000 -hrex
with different topologies (repl[0,1,2,3]/rest.tpr can be different)
Unlike PLUMED HREX, nstlist/dlb override is not necessary
